# Nitrate modulates stem cell dynamics by regulating *WUSCHEL* expression through cytokinins

**DOI:** 10.1101/200303

**Authors:** Benoit Landrein, Pau Formosa-Jordan, Alice Malivert, Christoph Schuster, Charles W. Melnyk, Weibing Yang, Colin Turnbull, Elliot M. Meyerowitz, James C.W. Locke, Henrik Jönsson

## Abstract

The shoot apical meristem (SAM) is responsible for the generation of all of the aerial parts of plants^1^. Given its critical role, dynamical changes in SAM activity should play a central role in the adaptation of plant architecture to the environment^2^. Using quantitative microscopy, grafting experiments and genetic perturbations, we connect the plant environment to the SAM, by describing the molecular mechanism by which cytokinins signal the level of nutrient availability to the SAM. We show that a systemic signal of cytokinin precursors^3^ mediates the adaptation of SAM size and organogenesis rate to the availability of mineral nutrients by modulating the expression of *WUSCHEL*, a key regulator of stem cell homeostasis^4^. In time-lapse experiments, we further show that this mechanism allows meristems to adapt to rapid changes in nitrate concentration, and thereby modulate their rate of organ production to the availability of mineral nutrients within a few days. Our work sheds new light on the role of the stem cell regulatory network, by showing that it does not only maintain meristem homeostasis but also allows plants to adapt to rapid changes in the environment.

---

Plants have evolved specific mechanisms to adapt their growth and physiology to the availability of mineral nutrients in their environment^5^. Various hormones such as auxin, abscisic acid, gibberellin and cytokinins have been shown to act in this process either locally or systemically^5^. Cytokinins in particular play an essential role in plant response to nitrate, where they act as second messengers^6^. For example, cytokinins promote lateral root development in areas rich in NO_3_ if the overall NO_3_ availability for the plant is low^7^. In the shoot, cytokinins have been shown to modulate key traits such as leaf size^8,9^ and branch number^10^ in response to nitrate.

Cytokinins have also been shown to be critical for the maintenance of stem cell homeostasis in the SAM. By modulating the expression of *WUSCHEL*, a homeodomain transcription factor expressed in the center of the meristem, cytokinins promote stem cell proliferation, thus controlling the size of the meristem and the rate of shoot organogenesis^11-14^. Using grafting experiments, a recent study showed that a systemic signal of a cytokinin precursor (trans-zeatin-riboside: tZR), travelling from root to shoot through xylem, could influence the size of the vegetative meristem^3^. However, it remains unclear whether cytokinin signalling can allow the SAM to respond to changes in nutrient levels in the environment. Here, we examined how a core stem cell regulator in the SAM dynamically responds to changes in mineral nutrient levels in the soil and whether systemic cytokinin signals can account for the dynamic adaptation of meristem function to nutrient levels. We used the inflorescence meristem of *Arabidopsis* as a model as this structure produces all of the flowers of the plant and is therefore a key target for crop improvement.

We first studied how fixed levels of nutrients affect meristem function, by growing plants on soil containing different levels of nutrients. To assess meristem function, we measured the size of the meristem, the plastochron ratio (a feature that is inversely proportional to the organogenesis rate of the SAM) and the number of flowers produced by the primary inflorescence (Methods). We observed that bigger, well-nourished plants had larger meristems and produced more flowers than smaller, more poorly nourished plants (Fig. 1a, Supplementary Fig. 1). We found a very close correlation between the weight of the rosette and the size of the meristem in individual plants (Fig.1b), showing that shoot development is coordinated when plants are grown under different nutritional conditions.

**Fig. 1.**
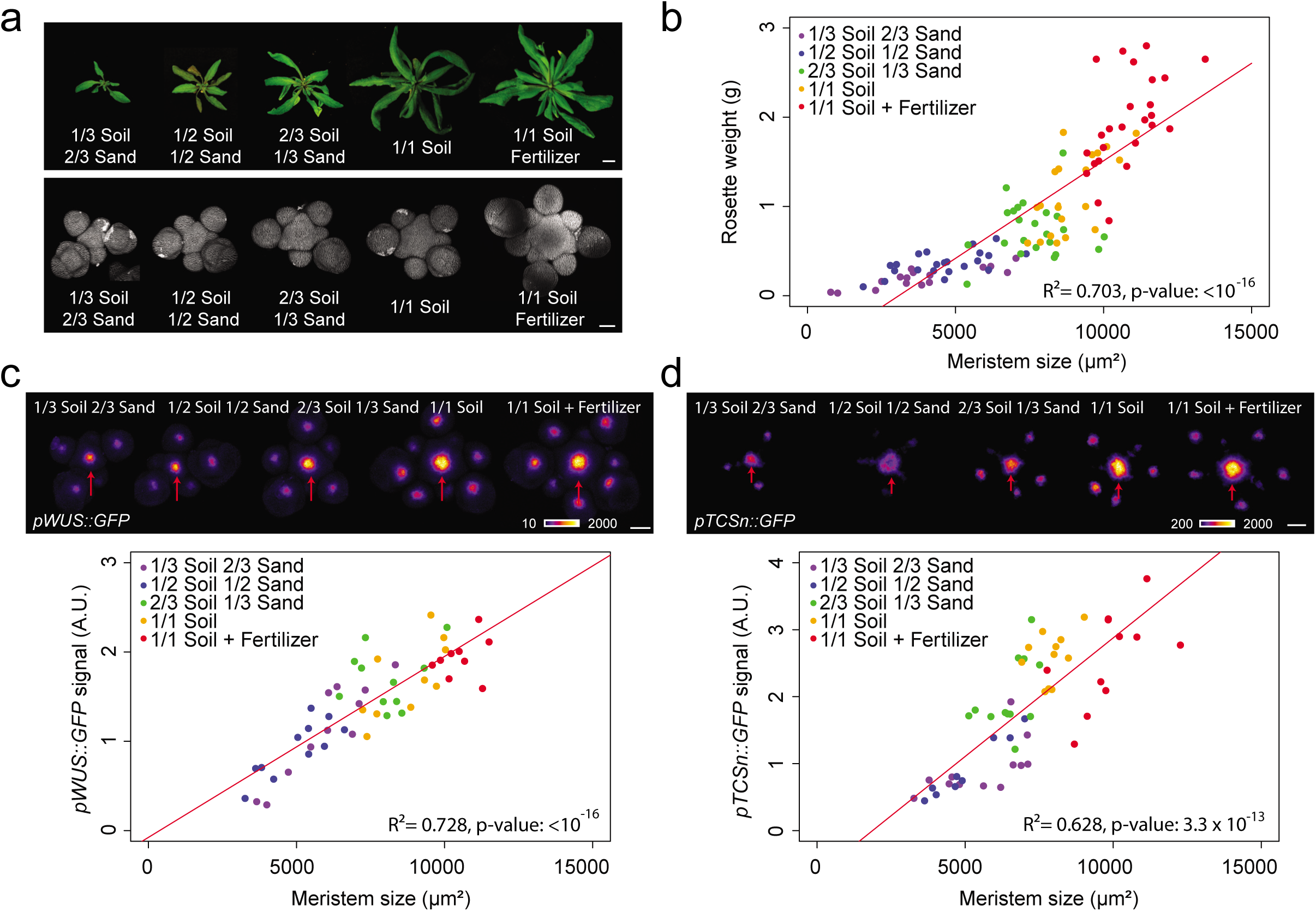
Plant nutritional status influences shoot meristem homeostasis. a. Morphology of the rosette (top, scale bar: 1 cm) and of the SAM (bottom, scale bar: 50 μm) of representative WT plants grown on soils of different nutritive quality. b. Correlation between rosette weight and SAM size of WT plants grown on soils of different nutritive quality (n=109, pool of 2 independent experiments). c. and d. *pWUS::GFP* (c) and *pTCSn::GFP* (d) expression in WT plants grown on soils of different nutritive quality (scale bars: 50 μm, n=55 (c) and n=57 (d)). Top panels show representative plants. Red arrows point to the center of the inflorescence meristem. The lower panels show total fluorescent signal (Methods) in the inflorescence meristem vs. meristem size. Data were fitted using linear models.

We next examined whether the observed changes in meristem size were linked to changes in stem cell homeostasis, by examining fluorescent reporters for the key meristem regulatory genes, *WUS* and *CLAVATA3* (*CLV3*). We developed a pipeline based on projections of 3D confocal stacks to automatically extract and analyse gene expression domains using a generalized exponential fit function (Methods, Supplementary Fig. 2). The intensity of the signal and the size of the domain of expression of *pWUS::GFP* were strongly correlated with the size of the SAM in the different growth conditions (Fig. 1c, Supplementary Fig. 3a and b), which was also confirmed using a translational fusion (Supplementary Fig. 3c). *CLV3* did not show such behaviour in all experimental repeats and only the size of its expression domain consistently correlated weakly with the size of the SAM (Supplementary Fig. 3d). Such an uncoupling between *WUS* and *CLV3* expression has been described in vegetative meristems under different light levels^15^ and suggests that these two genes could be differentially regulated by both nutrients and light.

As cytokinins modulate *WUS* and *CLV3* expression^11-13^, we examined the expression of *pTCSn::GFP*, a reporter of cytokinin response, when plants were grown with different levels of nutrients. Similarly to what was observed with *WUS,* the intensity of the signal and the size of the domain of expression of the *pTCSn::GFP* reporter correlated with the size of the meristem in the different growth conditions (Fig. 1d, Supplementary Fig. 3e and f), thus showing that plants growing with higher levels of nutrients exhibit higher levels of cytokinin signalling in the SAM. As altering cytokinin signalling is known to affect meristem homeostasis^16-21^, we further studied the phenotype in the meristem of various mutants of cytokinin metabolism and their response to mineral nutrition.

First, we looked at the phenotype of mutants altered in the successive steps of production of cytokinins^22^: *ipt3-1 ipt5-2 ipt7-1* (referred to here as *ipt3.5.7*) for the first step of biosynthesis, *log4-3 log7-1* (*log4.7*) and *log1-2 log3-2 log4-3 log7-1* (*log1.3.4.7*) for the second step of biosynthesis and for mutants for the conversion of cytokinins^23^: *cyp735a1-1 cyp735a2-1* (*cyp735a1.2*) and the degradation of cytokinins^16^: *ckx3-1 ckx5-2* (*ckx3.5*) in plants grown on soil supplied with fertilizer. Mutant shoots grew almost normally in this condition and only the *ipt3.5.7* and *cyp735a1.2* mutants showed a slight decrease in rosette weight (Fig. 2a and Supplementary Fig. 4a). However, meristem size and organogenesis rate were strongly affected by the mutations (Fig. 2b and c, Supplementary Fig. 4b-d). The *ipt3.5.7*, *log4.7, log1.3.4.7* and *cyp735a1.2* mutants, altered in various steps of tZ production, showed smaller inflorescence meristems that produced fewer organs. In contrast, the *ckx3.5* mutant, which displays higher levels of cytokinins, showed larger inflorescence meristems that produced more organs, as previously described^16^. We also quantified the size of the domain of expression of *WUS* by *in situ* hybridization (Supplementary Fig. 4e). We observed changes in the size of the *WUS* expression domain in all genotypes that correlated with the changes in meristem size we quantified, thus supporting the idea that cytokinin metabolism controls meristem function through the stem cell regulatory network.

**Fig. 2.**
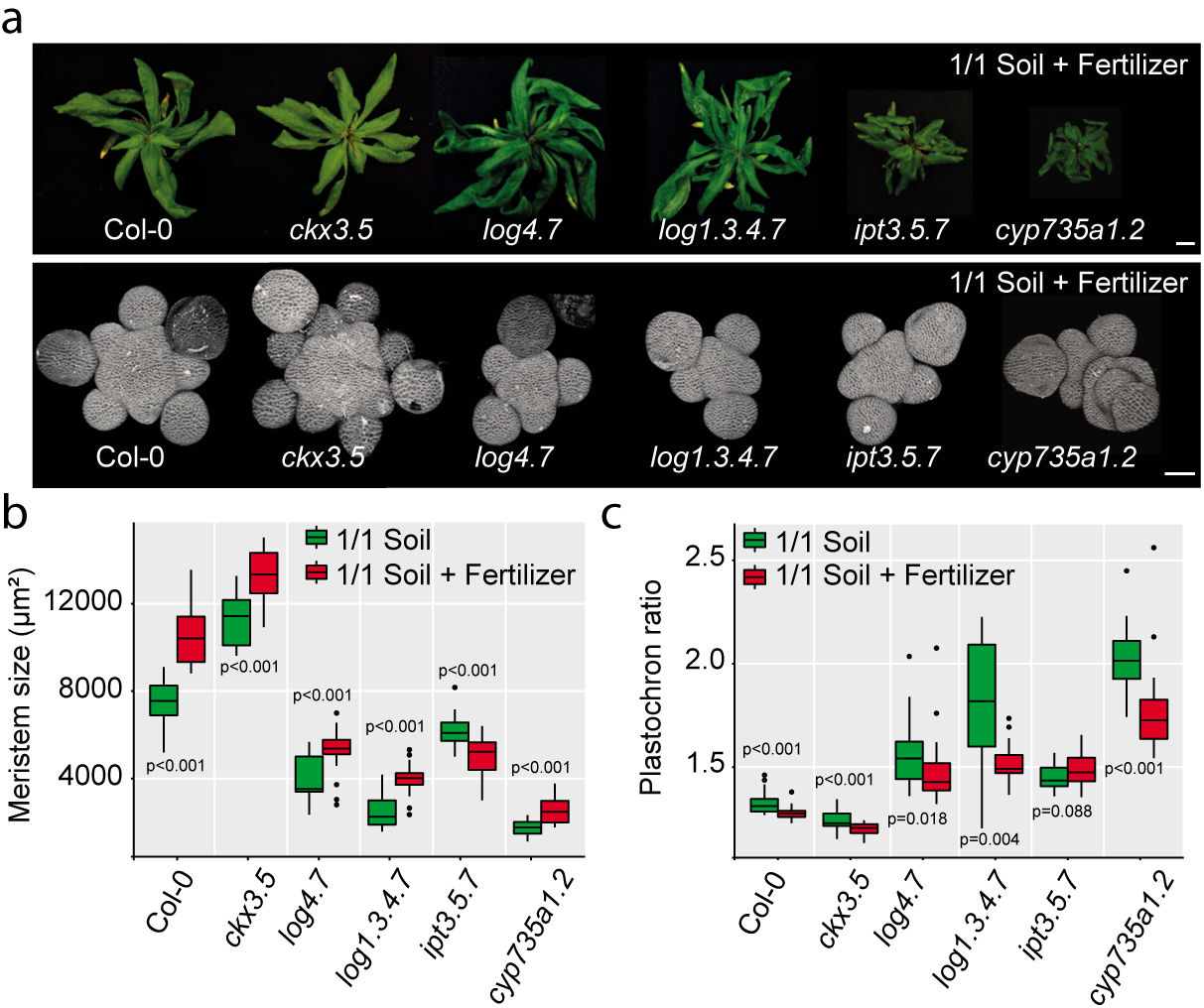
Cytokinins allow the adaptation of meristem function to plant nutritional status a. Morphology of the rosette (top, scale bar: 1 cm) and of the SAM (bottom, scale bar: 50 μm) of representative WT and CK-associated mutant plants. b. and c. Meristem size (b) and plastochron ratio (c) of WT and CK-associated mutant plants grown on soil without (green, g) or with fertilizer (red, r) (Col-0: n= 22 (g) and 25 (r), *ckx3.5*: n= 23 (g) and 27 (r), *log4.7*: n= 19 (g) and 24 (r), *log1.3.4.7*: n= 14 (g) and 25 (r), *ipt3.5.7*: n= 36 (g) and 20 (r), *cyp735a1.2*: n= 26 (g) and 20 (r), pool of 2 independent experiments). Data were compared using Student’s t-test.

We next looked at the effect of mineral nutrition on the phenotype of the mutants by comparing plants grown on soil with or without fertilizer. Of all mutants, only *ipt3.5.7* mutants did not respond to fertilizer with an increase in meristem size and organogenesis rate, as observed in WT. Instead, it displayed a surprising statistically significant decrease in meristem size (Fig. 2b and c). Although all the enzymes involved in cytokinin metabolism can modulate meristem homeostasis, *ISOPENTENYL-TRANSFERASE* enzymes (*IPT)*, which catalyse the first step of cytokinin production, appear to be the only ones necessary for the response of the meristem to changes in nutrient availability in the soil. This result is supported by the fact that nitrate, the main mineral nutrient, can directly modulate the expression of *IPT3* and to a lesser extent of *IPT5* in seedlings^24^. To further support that the response of meristems to mineral nutrients relies on cytokinin precursors produced by *IPT* enzymes, we used mass spectrometry to compare CK levels in inflorescences of plants grown without or with fertilizer. Despite technical variability in the mass spectrometry measurements, we observed a statistically significant increase in the levels of tZR precursors, the dephosphorylated products of *IPT* enzymes, in the two experimental replicates, further supporting the importance of these enzymes in the response of the meristem to nutrients (Supplementary Fig. 5f). We did not find significant changes in the levels of the active cytokinin tZ in the whole inflorescence but found a significant increase in the levels of the degradation products of tZ: tZROG (trans-zeatin ribose-O-glucose) and tZ7G (trans-zeatin-7-glucose). The lack of changes in tZ levels could be a result of tZ being a transient molecule mainly synthesized locally in the meristem, notably through the action of LOG4 and LOG7 enzymes^11-13^.

We further studied the spatial origin of the signal triggering the response of the meristem to mineral nutrition. Grafting experiments have previously shown that cytokinins can act either as local or as systemic signals in the control of root and shoot development^3,23,25^. We performed grafting experiments on our set of mutants to study whether a systemic signal is involved in the control of meristem function. Adding a WT root was able to rescue the phenotype in the meristem of *ipt3.5.7* and *cyp735a1.2* mutants but not of the *ckx3.5* and of the two *log* mutants (Fig. 3a and b, Supplementary Fig. 5a). Grafting a mutant root onto a WT scion led to a minor but statistically significant decrease in meristem size for *log1.3.4.7*, *ipt3.5.7* and *cyp735a1.2* mutant roots (Supplementary Fig. 5b-d). These experiments show that expressing *IPT* and *CYP735A* in either part of the plant is sufficient but not completely necessary for proper meristem function, but that *LOG* and *CKX* must act largely in the shoot to regulate meristem function. Our results are in agreement with the recent work by Osugi and colleagues, who showed that the phenotype in the vegetative meristem of the *ipt* triple mutant but not the *log* sextuple mutant could be complemented by grafting a WT root^3^. These results also make sense in light of the expression patterns of the different genes, as *LOG* and *CKX* enzymes are expressed in meristems^11,13,16^, while *IPTs* and *CYP735*a1-2 are expressed predominantly in the vascular tissues of the root and to a lesser extent of the shoot^19,26^ - only *IPT7* expression has been shown to be induced by *STM* in the vegetative meristem^27^.

**Fig. 3.**
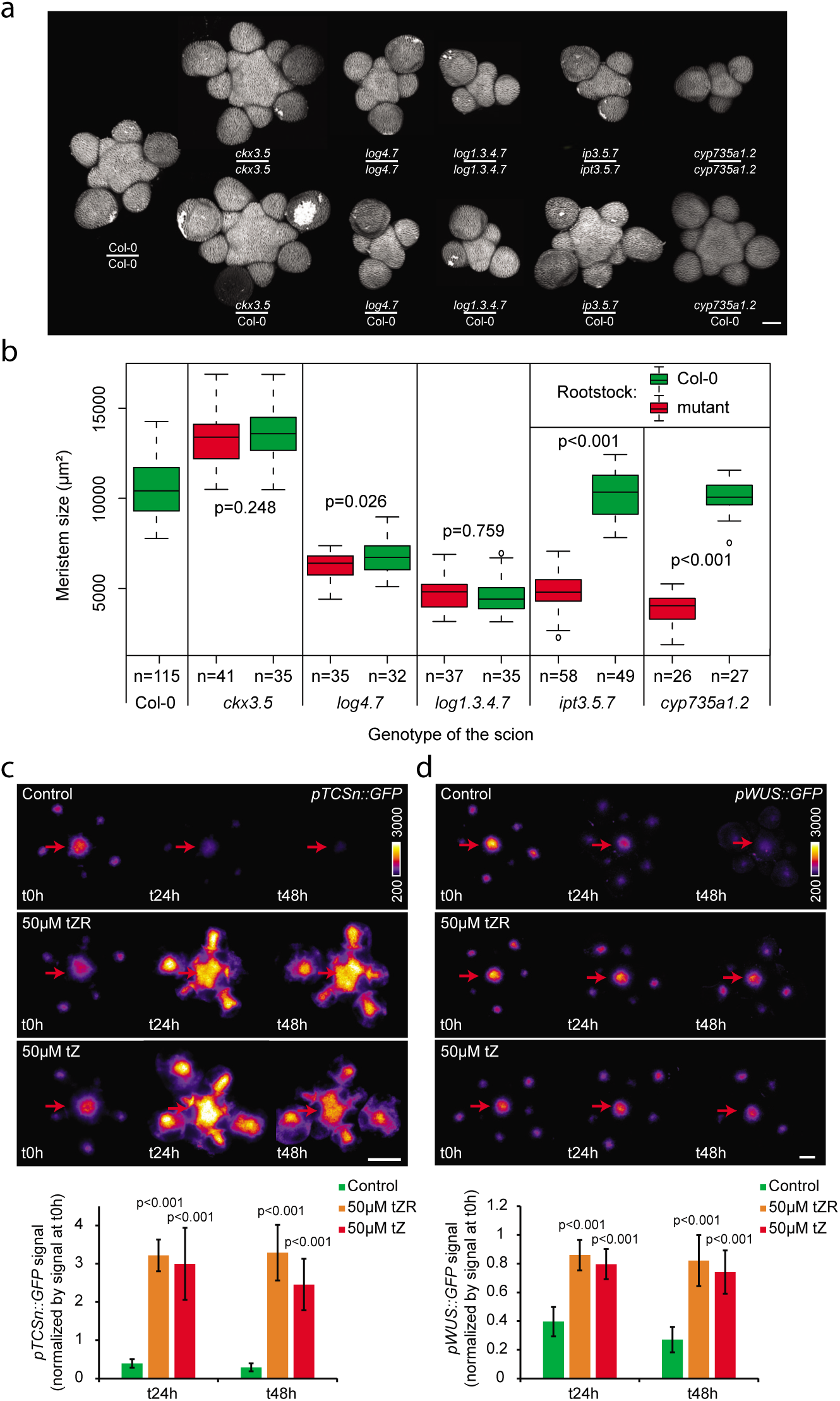
Cytokinin precursors act as long-range signals in the control of SAM homeostasis a. Representative meristems of WT and cytokinin-associated mutant plants self-grafted or grafted with a WT rootstock and grown on soil supplied with fertilizer (Scale bar: 50 μm). b. Meristem size of WT and cytokinin-associated mutant plants self-grafted or grafted with a WT rootstock and grown on soil supplied with fertilizer (pools of two independent experiments, the numbers of replicates are displayed in the figure). Data were compared using Student’s t-tests. c. and d. Effect of tZR and tZ application on the expression of *pTCSn::GFP* (c) and *pWUS::GFP* (d) in cut meristems grown *in vitro* (scale bars: 50 μm, pools of two independent experiments, *pTCSn::*GFP: n=11 for each condition, *pWUS::GFP*: n=12 for each condition). Top panels show representative plants. Red arrows point to the center of the inflorescence meristem. Lower panels show total florescent signal in the inflorescence meristem (Methods). Data were compared using Student’s t-test.

To analyse the dynamics between production of cytokinin precursors and modulation of meristem function, we applied tZR and tZ to meristems expressing *pTCSn::GFP* and *pWUS::GFP* cut from the plant and grown *in vitro* (see Methods). In the absence of added cytokinin, both *pTCSn::GFP* and *pWUS::GFP* signal strongly decreased over a 48h time course, suggesting that extrinsic cytokinin is needed to sustain meristem homeostasis. In contrast, adding 50 μM either of tZR or tZ to the medium upregulated *pTCSn::GFP* expression ectopically and allowed the maintenance of the *pWUS::GFP* signal in the centre of the meristem (Fig. 3c and d). Inducing *IPT3* in cut meristems in the absence of extrinsic cytokinin also maintained, at least partially, *WUS* expression (Supplementary Fig. 6). Together with the results from the grafting experiments, these data support a model where stem cell homeostasis is controlled by a systemic signal of cytokinin precursors (tZR) produced by *IPT* enzymes outside of the meristem, which are locally converted to active cytokinins by *LOG* enzymes in the shoot meristem.

Adding nitrate (NO_3_) to nitrogen-deficient plants leads to a rapid increase in cytokinin levels in the plant^3,24,28,29^. Given the rapid response of meristems to changes in cytokinin levels we observed *in vitro*, we looked to see if the meristem could dynamically respond to changes in nitrate concentration *in vivo.* We grew plants on sand, watering them with a nutritive solution containing a low concentration of nitrate (1.8 mM of NO_3_ as the only source of nitrogen) to induce a deficiency^10^. Once the plants bolted, we watered them with a nutritive solution containing different concentrations of NO_3_ (0 mM, 1.8 mM or 9mM) and quantitatively analysed the dynamics of response to this treatment. In the SAM, the treatment led to the dose-dependent induction of *pTCSn::GFP* and of *pWUS::GFP* expression within only one day and to an increase of meristem size and of the organogenesis rate within 2 to 3 days (Fig. 4a-c, Supplementary Fig. 7). We then checked whether this response relied on changes in cytokinin levels in the plant. In the root, we confirmed that the addition of nitrate to deficient plants led to a rapid, dose-dependent and transient induction of nitrate signalling (inferred from the level of expression of the nitrate responsive gene *NIA1*) and of *IPT3* and *5* (Supplementary Fig. 8a). By measuring the levels of cytokinin species by mass spectrometry, we also confirmed that the treatment led to a significant increase in the concentration of cytokinin precursors (tZR and/or tZRP depending on the tissues) and products of degradation (tZ7G, tZ9G, and/or tZROG depending on the tissues, Supplementary Fig. 8b). Finally, we analysed the meristem response in our set of mutants, three days after a nitrate treatment (9 mM NO_3_). Similar to what we observed on plants grown in soil, the response of *ipt3.5.7* mutant plants to nitrate in the meristem was strongly reduced, though statistically significant in one of the two experimental repeats, while response in the other mutant backgrounds was as in wild type (Fig. 4d, supplementary Fig. 9). In summary, our data show that the addition of nutrients leads to a response of the stem cell regulatory network within a day and that the rapid changes in meristem properties can be explained by the same cytokinin-based mechanism of signal propagation as we found for different fixed nutrient conditions (Fig. 2).

**Fig. 4.**
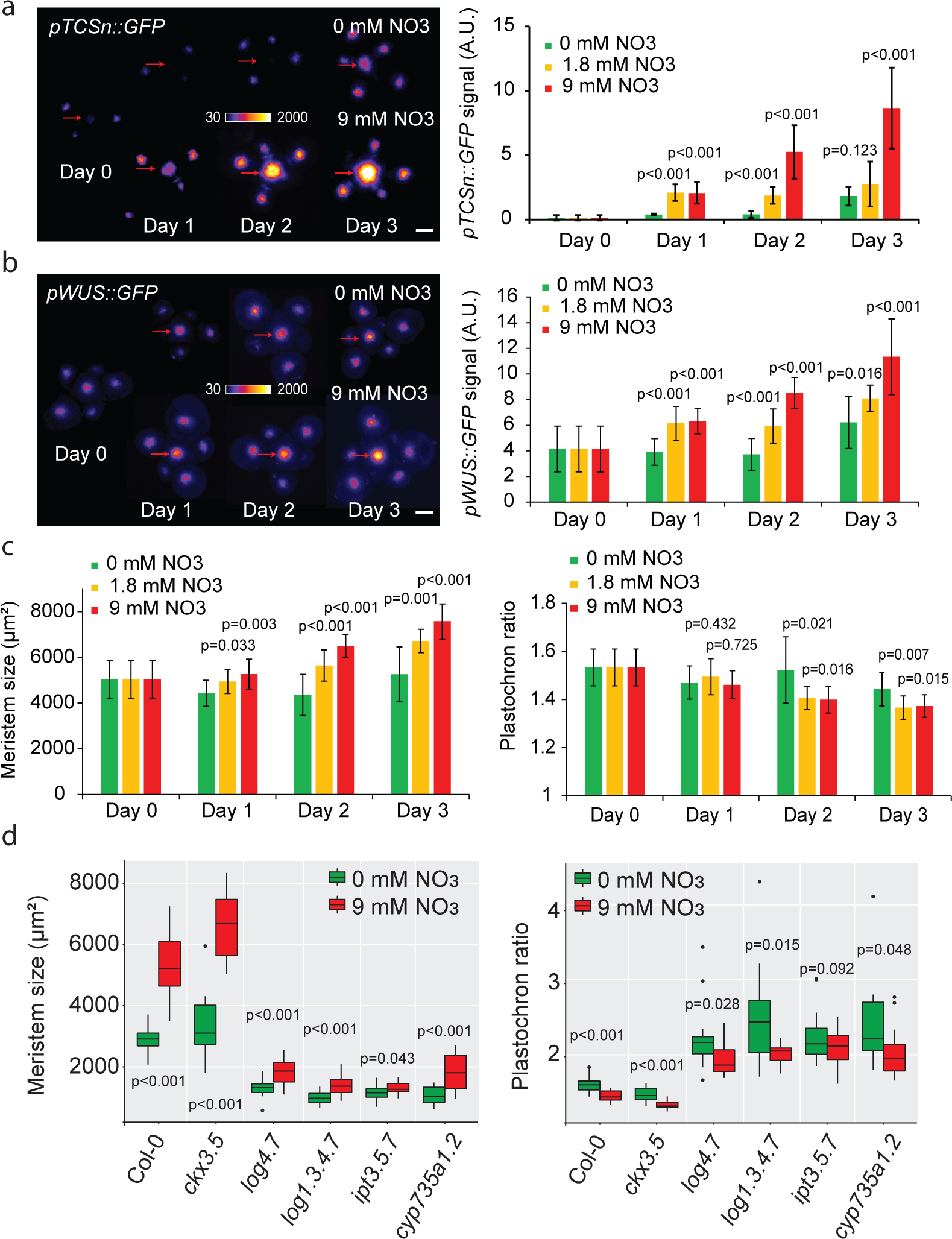
Nitrate modulates meristem homeostasis through *IPTs* a. to c. Effect of nitrate resupply on *pTCSn::GFP* expression (a), *pWUS::GFP* expression (b), and meristem size and plastochron ratio (c). The treatment was performed at day 0 and different meristems were dissected and imaged each day (Scale bars: 50 μm, n=8 to 12). (a and b) Left panels show representative plants, and right panels show quantification of total florescent signal (Methods). Red arrows point to the center of the inflorescence meristem. Data were compared using Student’s t-test. d. Meristem size and plastochron ratio of WT and cytokinin-associated mutants three days after treatment with a nutritive solution containing either 0 mM (green, g) or 9 mM of NO_3_ (red, r) (Col-0: n= 21 (g) and 24 (r), *ckx3.5*: n= 13 (g) and 12 (r), *log4.7*: n= 17 (g) and 18 (r), *log1.3.4.7*: n= 13 (g) and 15 (r), *ipt3.5.7*: n= 18 (g) and 19 (r), *cyp735a1.2*: n= 14 (g) and 14 (r)). Data were compared using Student’s t-test.

Taken together, our results show that the meristem can adapt the rate of shoot organogenesis to the availability of nitrate in the soil. Mechanistically, this phenomenon correlates with the ability of *WUS* to quantitatively respond to changes in the concentration of cytokinin precursors produced in the plant in response to variations in nitrate levels. As the main inflorescence of WT plants growing on soil only produced an average of 3.1 ± 0.4 flowers per day (n=16), the response of the SAM to changes in nitrate levels, which leads to significant changes in the rate of organ production in only two days, should be seen as very rapid in comparison to the pace of morphogenesis in this tissue. The timing of the response to nutrients is very similar to the response to induced perturbations of the core network^30^, in both cases causing expression domain changes in a day followed by changes in growth and size. Our findings thus expand the understanding of the function of the stem cell regulatory network in the SAM. They show that this network not only acts to maintain the integrity of the SAM during organogenesis, but also allows a very rapid adaptation of SAM function to a nutritional cue.

A recent study showed that vegetative meristems of seedlings germinating in the dark displayed reduced *WUS* expression, which was controlled by the activity of *CKX* enzymes^15^. Given the pleiotropic effects of cytokinins^6^, we can hypothesize that different environmental signals could be integrated through cytokinin signalling, and lead to different developmental responses in the meristem depending on whether they modulate, locally or globally, different aspects of cytokinin signalling.

Although it was developed in *Arabidopsis*, this model can in future be applied to crops, where cytokinin metabolism and action on stem cell regulation should be conserved. In rice, where *LOG* mutants were first characterized^18^, quantitative trait loci (QTL) for increased grain productivity have been mapped to genes involved in the regulation of cytokinins, including notably a CKX enzyme-encoding gene^31,32^. In maize, weak mutants of *FASCIATED-EAR3*, a receptor able to bind a *CLV3* homolog and involved in the control of *WUS* expression, also shows increased seed yield^33^. Our work, which provides a novel and integrative model based on nutrient availability, cytokinin metabolism and its effect on stem cell regulation and meristem function (Supplementary Fig. 10), allows a better characterization of the influence of mineral nutrients on plant architecture and could be used to better understand plant response to environmental inputs, and to develop new cultivars with increased yield.

## Author Contributions

B.L., C.W.M., E.M.M., J.C.W.L. and H.J. designed the experiments. B.L., A.M., C.W.M., C.S., W.Y. performed the experiments. B.L., P.F.-J., A.M., C.S. and C.T. analysed the data. B.L., E.M.M., J.C.W.L. and H.J. wrote the paper with inputs from all co-authors.

## Acknowledgments

This work is supported by the Gatsby Charitable Trust (through fellowship GAT3395/DAA to E.M.M., GAT3272/GLC to J.C.W.L. and GAT3395-PR4 for H.J.). E.M.M. also acknowledges support from the Howard Hughes Medical Institute and the Gordon and Betty Moore Foundation (through grant GBMF3406). P.F.-J. acknowledges a postdoctoral fellowship provided by the Herchel Smith Foundation. We thank Tanya Waldie and Maaike de Jong for sharing their knowledge on developmental plasticity and for help with the transient nitrate treatments, Jeremy Gruel for fruitful discussions on stem cell regulation, Paul Tarr for providing material and advice and Hugo Tavares for help with statistical analysis. We also thank Chillie Zeng, Mark Bennett and Rosa Lopez-Cobolla for their help during the preparation and the analysis of cytokinin species by mass spectrometry.

## Methods

### Plant Materials

Col-0 seeds were provided by the Salk stock center. The *pCLV3::dsRED* and *pWUS::GFP* marker lines were previously described^1,2^. The *pTCS::GFP* marker was provided by Teva Vernoux^3^ and the *pTCSn::GFP* was provided by Bruno Müller^4^.The *WUS-GFP* marker complementing the *wus* mutant phenotype was kindly provided by Jan Lohmann^5^. The triple *ipt* mutant *ipt3-1; ipt5-2; ipt7-1* (*ipt3.5.7*) was provided by Ottoline Leyser^6^. The *log4-3; log7-1* (*log4.7*), *log1-2; log3-2; log4-3; log7-1* (*log1.3.4.7*) and *cyp735a1-1; cyp735a2-1 (cyp735a1.2)* mutant seeds were kindly provided by Hitoshi Sakakibara and the RIKEN Institute^7^. The *ckx3-1* (SAIL_55_G12/CS870580); *ckx5-2* (Salk_117512C) double mutant has already been described^8^ and was generated by Paul Tarr.

### Growth conditions

For all plants grown on soil, batches of seeds were dispersed in pots of soil (Levington F2) and placed for 3 days in a 4°C room for stratification and then 7 days in a short day room (8h light) for germination. Seedlings were transferred in to separate pots containing different mixes of sand (Royal Horticultural Society: Silver Sand) and soil (Levington F2) and put into a constant light room (24 h light, temperature: 22°C, light intensity 160 μmol m^-2^ s^-1^). Plants were watered without or with 1/1/1 fertilizer (Vitafeed Standard). The experiments were carried out at the beginning of the flowering stage when the main inflorescence stem was of few centimetres in height.

For the nitrate resupply experiment, seedlings were germinated on soil and put in a short day room (8h light) for 7 days after stratification before being transferred into small individual pots of ½ sand (Leighton Buzzard sand from WBB Minerals) ½ Terra-green (Oil-Dri) pre-watered with 25 mL of an ATS-derived (*Arabidopsis* salts medium) nutritive solution containing 1.8 mM of NO_3_ prepared by adjusting the concentrations of the following components: 0.4 mM Ca(NO_3_)_2_, 1 mM KNO_3_, 4 mM KCl and 1.6 mM CaCl ^9,10^. Plants were put for 2 weeks in a short day room (8h light) before being transferred into a constant light room. Each plant was fed weekly with 10 mL of the same nutrient solution. The experiments were carried out at the beginning of the flowering stage when the main inflorescence stem was of few centimetres in height. At the beginning of the experiment (at day 0), plants were randomized, separated into three populations and watered with 25 mL of the nutritive solution containing either 0 mM of NO_3_ (by adjusting the concentrations of the following components: 0 mM Ca(NO_3_)_2_, 0 mM KNO_3_, 5 mM KCl and 2 mM CaCl_2_), 1.8 mM of NO_3_ (same solution as described earlier) or 9 mM of NO_3_ (normal ATS solution). Each day, 5-10 plants were used for analysis.

Within the course of the experiments, we realized that the tap water used for watering the plants contained non-negligible levels of NO_3_ (up to 0.7 mM in the summer), which could have moderately influenced our measurements and explain some of the differences observed between experimental repeats. Note however that within an experimental repeat, all genotypes or conditions were grown simultaneously and experienced the same growth conditions.

### Generation of the *p35S::XVE-IPT3-TFP* line

The coding sequence of *IPT3* was amplified from Col-0 cDNA (primers: F: TTTGGATCCTATGATCATGAAGATATCT-ATGGCTATG and R: AATGAATTCTCGCCACT-AGACACCGCGAC) and put by conventional cloning into a pENTR1Ad modified vector containing a synthetic sequence for a GS repeat linker peptide of 30 amino acids identical to the one described in^5^. Then, a LR recombination was performed using the *p35S::XVE-pENTR-R4-L1* plasmid described in^11^ and kindly provided by Ari-Pekka Mähönen, the *pENTR1Ad-IPT3-linker*, *TFP-pENTR-R2-L3* (Turquoise fluorescent protein) and the *pH7m34GW* destination vector coding for hygromycine resistance. The resulting plasmid was transformed into *Arabidopsis* plants carrying either both *pTCS::GFP* and *pCLV3::dsRED* or *pWUS::GFP* by flower dip transformation. Although we could see the effect of inducing *IPT3-TFP* in multiple independent insertions for each marker, we could only observe a very weak TFP signal by confocal microscopy.

### Image acquisition and time-lapse

Imaging was performed as follows: First, the main inflorescence meristem of plants at the beginning of the flowering stage was cut one to two centimetres from the tip, dissected under a binocular stereoscopic microscope to remove all the flowers older than stage 3 (as defined in^12^) and transferred to a box containing an apex culture medium (ACM) with vitamins as described in^13^. Meristems were imaged in water using a 20X long-distance water-dipping objective mounted either on a LSM700 or a LSM780 confocal microscope (Zeiss, Germany). Z-stacks of 1 or 2 μm spacing were taken. In some experiments, meristems were put in contact with a solution of 0.1 mg/mL of FM4-64 (Thermo-Fisher) for 10 minutes prior to imaging to dye the cell membranes. For the time-lapse experiment looking at the effect of extrinsic cytokinin, meristems were put in a box of ACM with vitamins and without or with 50μM of tZR or of tZ (Sigma) diluted from 50mM stock solutions. To prepare those stock solutions, tZR was dissolved in 10mM NaOH and tZ in DMSO. Meristems were kept in a constant light phytotron and covered with the same solution of liquid ACM for imaging. For the time-lapse experiment looking at the effect of an estradiol-based induction of *IPT3*, meristems were put in a box of ACM with vitamins and with 5 μM of β-estradiol (Sigma) or with the equivalent volume of DMSO as a control. Again, meristems were kept in a constant light phytotron and covered with the same liquid ACM solution for imaging.

### Image Analysis

Confocal stacks were analysed using the ImageJ software (https://fiji.sc/). Z-projections (maximum intensity) of the stacks were performed to analyse meristems dyed with FM4-64. Z-projections (sum slices) of the stacks were performed to analyse meristems expressing *GFP* or its derivatives and the “fire” lookup table was used to represent the signal. Meristem size and plastochron ratio were measured as described in^14^. Note that as the plastochron ratio is the average area between successive primordia, this parameter is more variable for meristems producing few organs as it is averaged on a smaller number of measurements. The expression level of *pTCSn::GFP* and the expression level and domain size of the *pWUS::GFP*, *WUS-GFP* and *pCLV3::dsRED* were analysed using a Matlab pipeline (Mathworks Inc., Natick, MA) specifically developed for this study (see below).

### Characterization of gene expression domains through automated image analysis

In order to characterize quantitatively the *WUS*-*GFP*, *pWUS::GFP*, *pTCSn::GFP* and *pCLV3::dsRED* domains in meristems, we developed an automatic pipeline with custom-made code written in Matlab. The code performs two basic tasks: first of all, it automatically identifies the fluorescent domain in the inflorescence meristem. Second, it computes the size and the intensity of such a domain. The domain identification is performed on Gaussian filtered (σ=5μm) maximal intensity projection of the fluorescence image under study, while the size and intensity of the domain calculation is performed on a Gaussian filtered (σ=5μm) sum slices projection. In both tasks, the Gaussian filter is used to smooth out the fluorescence images so that we neutralize the intracellular localization (ER or nucleus) of the fluorescent markers. This produces a more continuous fluorescence domain, which facilitates the automated extraction of its main features. Below, we explain in further detail how this pipeline was structured and used.

The automatically identification of the center of *WUS* and *CLV3* expression domains is performed through an Otsu thresholding over the corresponding z-projected and Gaussian-filtered fluorescence images. As floral primordia also express the reporters, it is assumed that the expression domain of the inflorescence meristem is the one that is the closest to the center of each image. Then, the centroid, ***r*_*0*_**, of this domain is found and a circular region *R* of radius *ρ*_*R*_ =40 μm is defined that should contain the fluorescence domain under study in the meristem. In the cases where the expression domain has failed to be automatically detected – e.g., in those images where the meristem was not well centered – we have manually specified the center of the inflorescence meristem. *i+1*

After domain detection, the size of fluorescence domains is determined. To do so, the radially symmetric decay of the fluorescence in the *R* regions of the corresponding z-projected and Gaussian filtered images is characterized. Specifically, a characteristic length *L*_*0*_ of such decay in each *R* region is extracted, and we interpret this as the fluorescence characteristic domain size. To extract *L*_*0*_, the mean fluorescence intensity within different concentric subregions within region *R* around the centroid ***r*_*0*_** is computed; the most central subregion is a circle of radius *dρ*=1 μm, and the other subregions are 39 concentric i-toroids (*i=1,…39*) filling the *R* region; the *i-*th toroid is defined by those pixels whose (x,y) coordinates fulfill the relation *ρ_i_ ^2^<(x-x_0_)^2^+(y-y_0_)^2^ ≤ ρ_i+2_^2^*, *x*_*0*_ and *y*_*0*_ being the coordinates of the centroid ***r*_*0*_** and where *ρ*_*i*_ *=idρ*. Then, the characteristic length *L*_*0*_ is extracted by fitting the fluorescence profile to an exponential function of the form ^15^

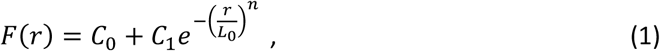

where *C*_*0*_ corresponds to the mean autofluorescence levels of the sum slices projection, *C*_*1*_ is the height of the fluorescent peak, and *n* is an exponent accounting for the radial decay of the fluorescent domain. *C*_*0*_, *C*_*1*_ and *n* are also determined in the fit.

The total fluorescence of the selected domains (referred as signal), *M*_*R*_, is calculated by

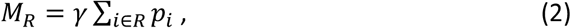

where *p*_*i*_ is the pixel intensity levels in the *i*-th pixel of the sum slices projection, and *γ* is the resolution in microns squares of the image under study. The *γ* factor corrects the differences in intensity levels between images that have different resolution. Then, the inferred autofluorescence, *B*_*R*_, is subtracted from the summed fluorescence in the selected domain, so that the intensity used in the analysis reads

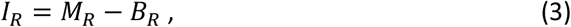

where *B*_*R*_ =*C*_*0*_ *A*_*R*_, being *A*_*R*_ the area of the *R* region.

We have checked that when fixing the *n* exponent as the average exponent for all the meristems in a given experiment, we obtained equivalent results for the characteristic length and total fluorescence to the case where the *n* exponent was a free parameter.

The code can be found following this link: https://gitlab.com/slcu/teamHJ/pau/RegionsAnalysis(v0.1)

### RNA extraction and qPCR

Three biological replicates of three roots each were used in each time-point. Roots were quickly washed in water to remove the sand and terra-green and frozen in liquid nitrogen before being manually ground. Total RNA was extracted using the RNA plus mini kit (Qiagen). First strand cDNA was then synthesized using the Transcriptor First strand cDNA kit (v.6 Roche). Real-time quantitative PCR was performed using double-strand DNA-specific dye SYBR Green (Applied Biosystems) in a LightCycler 480 II (Roche). positive replicates and one negative (using as template a product of the cDNA transcription made without the transcriptase) were performed for each reaction. Expression of *UBQ10*, *NIA1*, *IPT3* and *IPT5* was assessed using the primers described in the following table. Expression data of *NIA1*, *IPT3* and *IPT5* were obtained by normalizing for each replicate the ½ CT of the corresponding gene by the one of *UBQ10*.

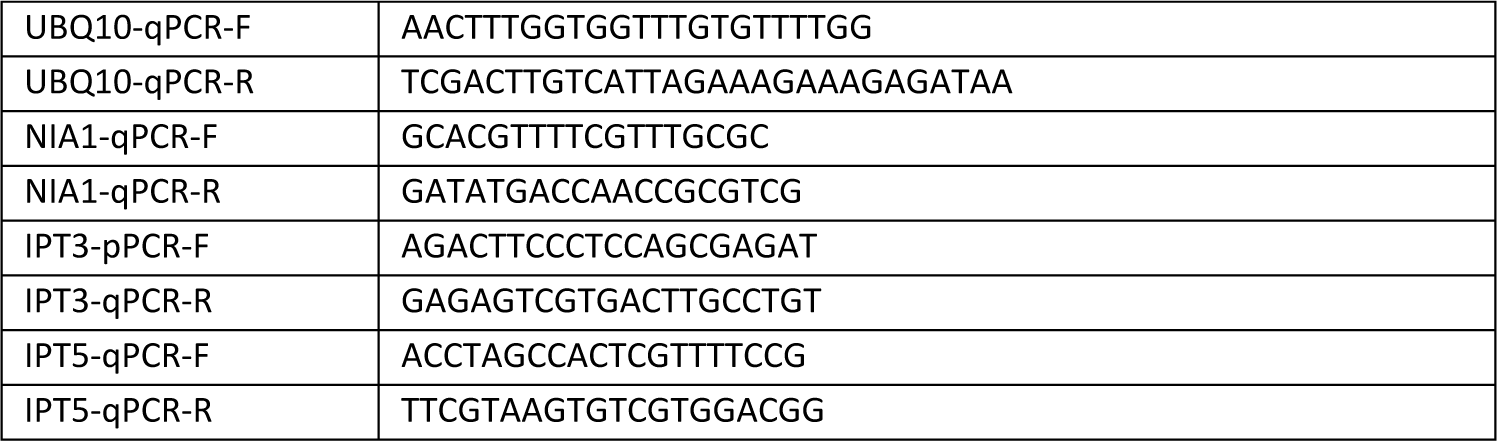

### Cytokinin analysis by mass spectrometry

Three replicates of three roots, shoots or inflorescences each were used in each condition. Tissues were frozen in liquid nitrogen, ground manually and an aliquot of 100 to 400 mg was taken and weighed. Cytokinins were then extracted and measured by liquid chromatography - mass spectrometry (LC-MS) following the protocol described in ^16^, with the following modifications: the mass spectrometer was an Applied Biosystems QTrap 6500 system, and the initial mobile phase was 5% acetonitrile in 10 mM ammonium formate (pH3.4).

### Grafting

*Arabidopsis* micro-grafting was performed on 7-day-old seedlings according to the protocol described in^17^. Grafted seedlings were transferred to soil 7 days after grafting, watered with 1/1/1 fertilizer and put in a constant light room until bolting stage.

### *In-situ* Hybridization

Full length *WUS* cDNA was amplified by PCR using specific primers and ligated into a pGEM-T Easy vector (Promega). Transcription was performed using the DIG RNA labelling kit (Roche). Embedding of the samples, sectioning (8μm sections) and *in situ* hybridization were performed following the protocol described in (http://www.its.caltech.edu/∼plantlab/protocols/insitu.pdf) with the addition of 4% polyvinyl alcohol (PVA) to the nitro blue tetrazolium-bromochloroindolyl phosphate (NBT-BCIP) staining solution. Sections were captured with a 20x lens using an Axioimager microscope (Zeiss) with fully opened diaphragm at 2048 x 2048 picture size. Quantification of the *WUS* expression area was described previously^18^. The threshold was determined by the mean intensity of the unstained tissue of the same section minus four standard deviations. Three consecutive sections that showed the strongest staining signal were analysed per meristem and the expression area was averaged. For *ipt3,5,7* and *cyp735a* the staining signal of two consecutive sections was quantified and averaged, since the *WUS* signal was only detectable in two sections. Image analysis was performed using the ImageJ software.

### Statistical analysis

Statistical analyses were performed using either Microsoft Excel or R software (https://www.R-project.org). Mean or mode values (for boxplots) were shown with error bars corresponding to the standard deviation for all experiments except for the analysis of the levels of cytokinin species by mass spectrometry where mean was shown with error bars corresponding to the standard error due to the low number of experimental repeats. The number of biological repeats (usually the number of meristems) is displayed in the legend of each figure or within the figure itself. When independent experiments could not be pooled, one independent experiment was shown in the main figures and one in the supplementary material. When applicable, data were fitted using either linear or exponential models depending on the shape of the cloud of points and on the value of the coefficient of determination R^2^ and of the p-value of an F-test on the fit. Within the same experiment, the same type of fit was used when comparing data from different genotypes. When Student tests were performed to compare time points and genotypes. The measurements were assumed to have two-tailed distributions and unequal variances and were considered as significantly different when the p-value was lower than 0.05.

